# *In silico* discovery of small molecules for efficient stem cell differentiation into definitive endoderm

**DOI:** 10.1101/2021.10.06.463418

**Authors:** Gherman Novakovsky, Shugo Sasaki, Oriol Fornes, Meltem E. Omur, Helen Huang, Nathaniel Lim, Artem Cherkasov, Paul Pavlidis, Sara Mostafavi, Francis C. Lynn, Wyeth W. Wasserman

## Abstract

Improving methods for human embryonic stem cell differentiation represents a challenge in modern regenerative medicine research. Using drug repurposing approaches, we discover small molecules that regulate the formation of definitive endoderm. Among them are inhibitors of known processes involved in endoderm differentiation (mTOR, PI3K, and JNK pathways) and a new compound, with an unknown mechanism of action, capable of inducing endoderm formation in the absence of growth factors in the media. Optimization of the classical protocol by including this compound achieves the same differentiation efficiency with a 90% cost reduction. The gene expression profile induced by the compound suggests that it is an inhibitor of the MYC pathway. The proposed *in silico* procedure for candidate molecule selection has broad potential for improving stem cell differentiation protocols.

## Introduction

Over 400 million people worldwide live with diabetes, a chronic disease caused by insulin deficiency or resistance. Common treatments for diabetes include dietary regulation and insulin injections, however, a promising emerging approach involves the use of human embryonic stem cell (hESC)-derived insulin-producing cells to replace patient cells that are lost or dysfunctional (reviewed in (Migliorini, Nostro, and Sneddon 2021)). Current protocols for generating insulin-producing cells from hESC mimic the specification of pancreatic β cells during development by differentiating hESCs into primitive streak (PS) (Davis et al. 2008), an intermediate fate branch point (Davis et al. 2008), and then definitive endoderm (DE), a crucial pancreatic precursor (Zorn and Wells 2009; Lee and Chung 2011; Mahaddalkar et al. 2020) (**Fig. S1A**). Activation of the TGFβ and WNT pathways through extrinsic signals, and the inhibition of the PI3K/mTOR pathway, lead hESCs to the PS branch point. PS cells form the mesendoderm, a short-term *in vitro* precursor of endoderm and mesoderm (ME) lineages (Rodaway and Patient 2001). Prolonged treatment of mesendoderm cells with WNT and BMP inducers leads to ME lineage, while downregulation of WNT/BMP signaling and activation of the TGFβ pathway induce DE formation. These processes are well studied with well-known gene markers (**Table S1**). Improving differentiation protocols for DE from hESCs remains a focus of much research, with some protocols using CHIR99021 (henceforth CHIR), a WNT inducer, and the growth factor Activin A (AA), a TGF β inducer (Loh et al. 2014; Naujok, Diekmann, and Lenzen 2014) (**Fig. S1B**).

The complex manufacturing process and high cost of growth factors like AA (750-2,500 USD per 100 μg (“Human Activin A Recombinant Protein” n.d., “Human Recombinant Activin A” n.d.)) makes current DE differentiation protocols expensive. Small molecules are ideal replacements: they are more stable, easier to store, allow for greater specificity, and have greater activity and reproducibility (reviewed in (Pan and Liu 2019)). Several studies have shown that small molecules can successfully induce differentiation (Williams et al. 2007), even in the absence of growth factors (Korostylev et al. 2017; Borowiak et al. 2009). However, small molecules rarely act on one single target, which can cause undesired off-target effects (De et al. 2017), and their discovery, for instance by high-throughput screening, is an expensive and time-consuming process (Waring et al. 2015).

Drug repurposing is an alternative method for identifying compounds with desired properties (reviewed in (Pushpakom et al. 2019)). The Connectivity Map (CMap) is a catalog of induced gene expression signatures for thousands of compounds (Lamb et al. 2006). It provides a platform for identifying potential inducers/inhibitors of a process of interest based on the similarity of the expression signatures of the process and those of compounds in the database. The first release of CMap included 6,100 gene expression profiles from 1,309 unique compounds tested in 5 different cell lines. The application of CMap for drug repositioning has been demonstrated in different fields (J. Liu et al. 2015; M. Zhang et al. 2015; Dyle et al. 2014; Farooq et al. 2009; Dudley et al. 2011), including the identification of small molecules inducing osteoblast differentiation (Brum et al. 2015, 2018). Recent studies increasingly utilize the Library of Integrated Network-based Cellular Signatures (LINCS) database, an expansion of the CMap catalog (the second release) including >500,000 gene expression signatures from the screening of >20,000 small molecules across 99 different cell lines (Subramanian et al. 2017).

Here, using DE differentiation as a model developmental process, we describe three *in silico* screening approaches to discover small molecules that can be used for directed differentiation. To accomplish this, we used transcriptomic profiles of AA-based DE differentiation or pathway and TF target enrichment to query the CMap/LINCS catalogs and identify candidate AA replacements. We tested the ability of a subset of these candidates to drive endoderm differentiation from hESCs, including a new compound that might regulate DE formation through altering MYC activity. This approach presents an efficient alternate means to optimizing stem cell differentiation protocols.

## Results

### Bulk RNA-seq analysis of public datasets reveals definitive endoderm-specific TFs and pathways

To identify potential inducers of endoderm differentiation, we built a profile of gene expression changes of hESC differentiation into DE (**Fig. S1A**). We analyzed two publicly available bulk RNA-seq time-series datasets that capture gene expression changes in hESCs after the addition of AA to the media (**Fig. S2A**): GSE75748 (Chu et al. 2016) (H1 hESCs; time points 0, 12, 24, 36, 72, and 96h) and GSE109658 (Lu et al. 2018) (H9 hESCs; time points 0, 24, 48, 72 and 96h). We then performed differential expression analysis at 0 *(i.e.* pluripotent stage) and 96h (*i.e.* DE stage), and compared the list of differentially expressed genes between the two datasets (**Methods**). Both up- and down-regulated genes significantly overlapped between the two datasets (hypergeometric test, *p*-value = 0), with 735 and 433 genes, respectively (**Table S2**). The analysis confirmed not only the agreement between the two datasets (**Fig. S2B**), but also highlighted groups of genes with similar and expected expression patterns (**Table S1**). For instance, the pluripotency markers *POU5F1*, *SOX2*, and *NANOG* showed a steady decrease of expression over time. In contrast, expression of mesendoderm markers, such as *TBXT*, *FGF4*, or *CDX1*, peaked at 24h, declining shortly after, while the expression of endoderm markers including *SOX17*, *GATA6*, and *EOMES* increased over time (**Fig. 1A**). Notably, the number of differentially expressed genes at 96h was in the order of thousands, which is consistent with the significant changes that cells undergo in response to AA treatment (Chu et al. 2016) (**Fig. S2C**).

**Fig. 1.**
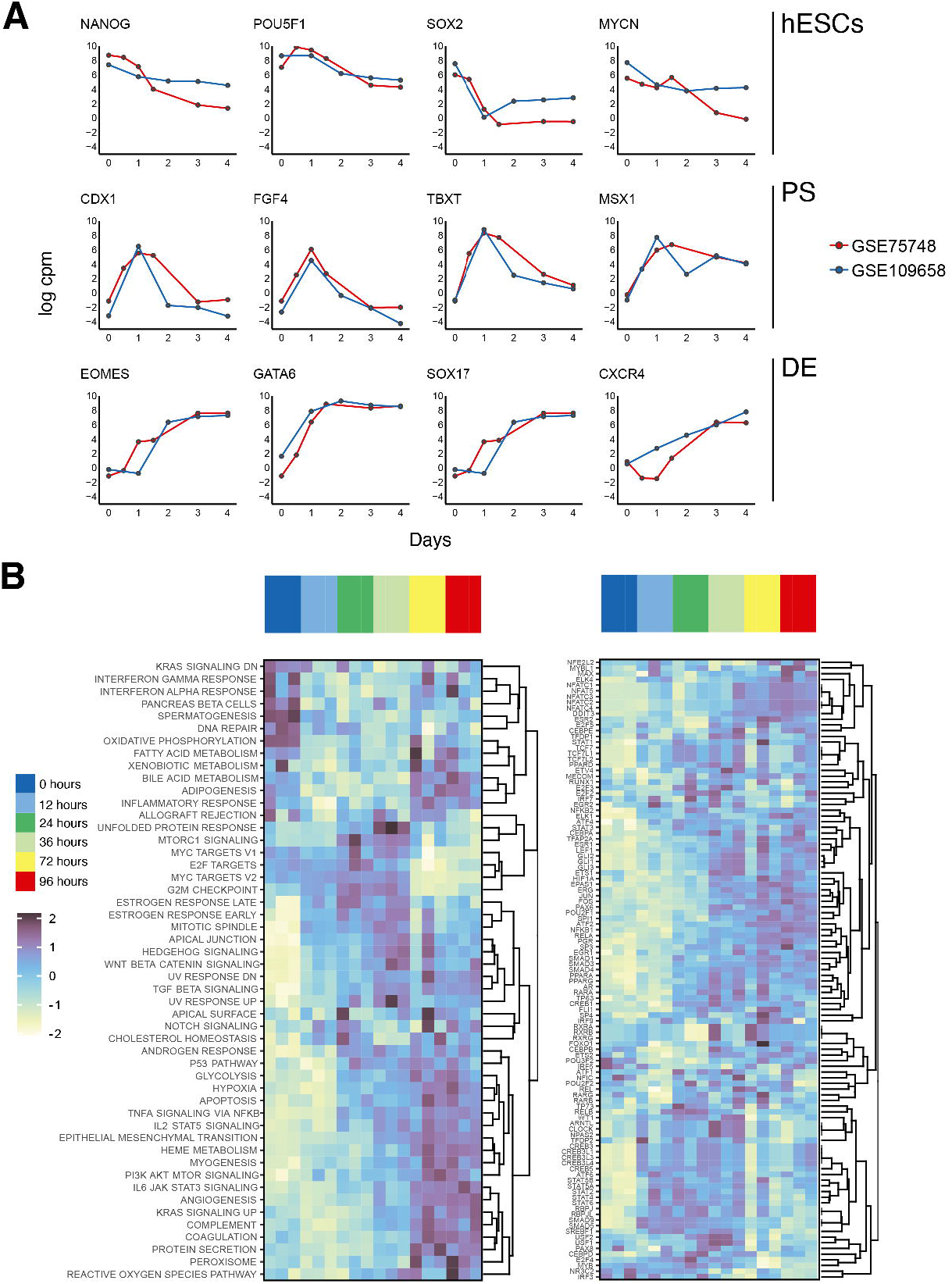
Analysis of public DE differentiation datasets. **A** - the expression trend of key genes involved in pluripotency and mesendoderm/endoderm development; shows consistency among public datasets; **B** - heatmaps with activation values of pathways and transcription factors along differentiation timeline respectively; clustering is based on GSE75748 values.

Next, we explored pathway activities at each time point using single sample gene-set enrichment scores (Hänzelmann, Castelo, and Guinney 2013; Barbie et al. 2009) (**Methods**; **Figs. 1B** and **S3**, left panels). For ease of interpretation, estimated activities of Hallmark pathways from the Molecular Signatures Database (MSigDB) (Liberzon et al. 2015) were clustered. DNA repair and spermatogenesis pathways were significantly upregulated in early stages (**Table S3**), consistent with previous reports linking cell cycle-related pathways with pluripotency (Pauklin and Vallier 2013; Chappell and Dalton 2013; Hsu et al. 2019). Similarly, MYC and E2F targets showed a strong initial activity that became negligible in later time points. The following pathways were activated at later time points: p53, apoptosis, hypoxia, epithelial-mesenchymal transition (EMT), PI3K/AKT/mTOR, TNFα via NF-κB, IL-2/STAT5 signaling, WNT, and TGFβ. All identified pathways were significant (**Table S3**). The activation of several pathways was consistent with expectations, for instance TGFβ pathway as it is induced by AA, p53 pathway which has been implicated in DE differentiation (Wang et al. 2017) and apoptosis pathway as it shares similarities with the differentiation processes (Launay et al. 2005; Lanneau et al. 2007). On the other hand, activation of the PI3K/AKT/mTOR pathway was surprising, as it has been shown to promote neuroectoderm differentiation (Yu and Cui 2016), and the roles in DE differentiation of NF-κB and IL-2/STAT5 pathways were unclear.

Using the same procedure as above, we assessed TF activity given the enrichment of their respective targets (**Figs. 1B** and **S3**, right pannels). POU3F and NFI, among others (see **Table S3** for full list) showed enhanced activity in early time points, which is consistent with their roles in neural plate development, possibly indicating the “readiness” of stem cells to adopt a neuroectoderm fate while awaiting an external signal (Iwafuchi-Doi et al. 2011; Mason et al. 2009; Boheler 2009). In the subsequent 48h, different TFs became active. Interestingly, receptor-regulated SMADs displayed different patterns of activity. It is known that SMAD5 and 9, which are involved in BMP signaling (Retting et al. 2009), are crucial for ME development, while SMAD2 and 3, which mediate Activin/Nodal signaling (Fei et al. 2010), promote hESC to endoderm differentiation. Therefore, the observed increase in SMAD5 and 9 activity around mesendoderm formation and their subsequent steady decline was not surprising, while SMAD3 and its co-regulator SMAD4 exhibited a stable increase. Notably, SMAD1, despite being BMP-dependent like SMAD5 and 9, displayed a similar activation pattern to SMAD3 and 4. The activity of TFs GLI1, 2, and 3 increased steadily over time, in agreement with the important role of the Hedgehog signalling pathway in endoderm development (Deol et al. 2017). We also observed an increase of HIF1A activity, which is a hypoxia-inducible factor. The presence of JUN factor activity was interesting, as we could not find prior reports linking it to the formation of endoderm lineage (**Fig. S3,** right).

Taken together, the analysis of the bulk RNA-seq data highlighted the importance of TGFβ, hypoxia, mTOR, and other pathways, as well as the TFs SMAD3 and 4 in the differentiation process.

### Single-cell RNA-seq analysis of the primitive streak branch point identifies potential regulators of definitive endoderm fate

Cell differentiation is a complex, heterogeneous and continuous process; its analysis at single-cell resolution could potentially reveal novel TFs regulating the differentiation process or guiding cell fate decisions at branch points, as well as new intermediate cell types. We therefore performed scRNA-seq of the PS branch point (**Fig. 2A**). We profiled at 36 and 72h post-induction with AA, and at 72h post-induction with CHIR, which induces ME differentiation (**Methods**). After data pre-processing and clustering, cells were annotated into different developmental stages (hESC, PS, ME, and DE) based on known gene markers whose expression correlated well with the sampled time points (**Fig. 2B**; **Table S1**). Then, we identified differentially expressed TFs in DE, ME, and PS (**Methods**). We hypothesized that TFs inducing DE lineage would be differentially expressed in both PS (the branch point) and DE. Similarly, common differentially expressed TFs in PS and ME would induce ME lineage. The set of DE inducers included TFs such as EOMES (Teo et al. 2011), GATA6 (Fisher et al. 2017), GSC (Yasunaga et al. 2005), HHEX (Martinez Barbera et al. 2000), LHX1 and OTX2 (Costello et al. 2015), PITX2 (Faucourt et al. 2001), TGIF1 (Powers et al. 2010), and ZIC2 (Warr et al. 2008), whereas FOXH1 (Slagle, Aoki, and Burdine 2011), MSX1 (Hara and Ide 1997), SP5 (Weidinger et al. 2005), TBXT (Faial et al. 2015), and TBX6 (Chapman et al. 2003) were among the set of ME inducers (see **Table S4** for the full sets). Pseudotime analysis revealed that both groups of genes became strongly upregulated through the course of differentiation at each respective stage, supporting the role of these TFs in regulating lineage specification (**Figs. 2C** and **D**).

**Fig. 2.**
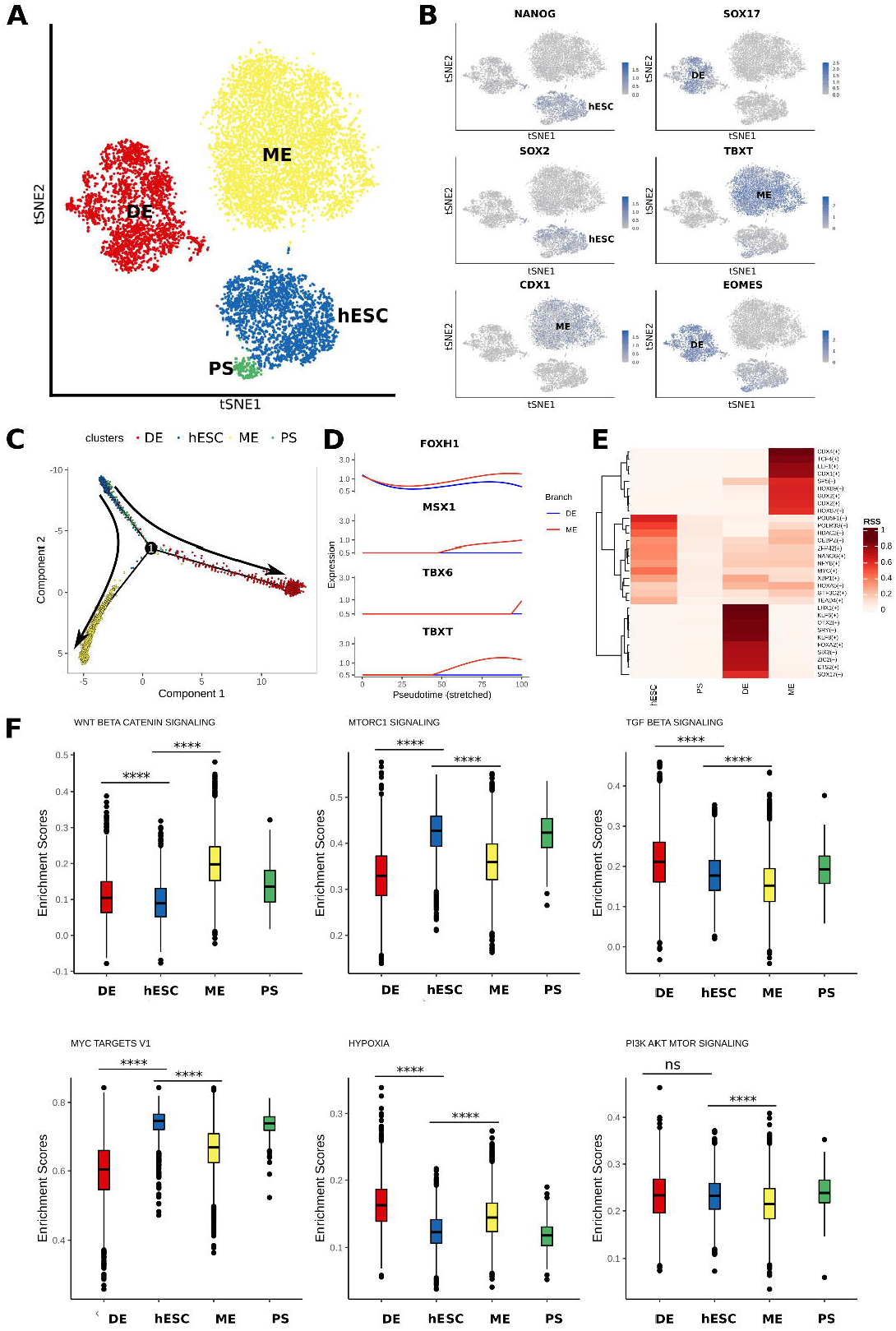
Single-cell RNA-seq analysis. **A** - tSNE plot of the analyzed dataset; major cell clusters are labeled with the corresponding cell type names; **B** - distribution of the expression values of the marker genes, which were used to annotate the clusters; **C** - pseudotime ordering of the cells shows the expected bifurcation point with DE and ME branches; **D** - expression trends of the key mesoderm markers along the differentiation axis. The absence of the “spike” that is present in bulk data (**Fig. 1A**) can be explained by low depth and resolution of the single-cell data; **E** - the RSS score heatmap of the most informative TFs for each of the cell types; **F** - the distribution of GSEA scores per single-cell for the key developmental pathways. The stars (****) highlight the significance of the change with respect to hESC (Wilcoxon test).

Next, we identified“active” TFs during the course of differentiation, using an approach called SCENIC (Aibar et al. 2017) (**Methods**). Overall, SCENIC results were in agreement with those of the bulk RNA-seq analysis. Consistent with previous reports, SCENIC identified endoderm-related markers, including LHX1 and KLF6/8 (Chu et al. 2016), OTX2 and FOXA2 (Fu et al. 2011), SOX17 (Qu et al. 2008), EOMES and ETS1/2 (Mohammadnia et al. 2016), FOXQ1 (Shiraki, Ogaki, and Kume 2014), and SIX3 (Jing Wen et al. 2017). It also revealed potentially new markers such as RREB1, CUX1, IRF9, DDIT3, TFAP2C, PBX3, JUN, and JUND (**Fig. 2E**). Notably, RREB1 (Lee et al. 2012) and CUX1 (Ripka et al. 2010) are important for the development of the primitive gut tube and pancreas, while DDIT3 is a known apoptosis inducer (Papathanasiou et al. 1991), and TFAP2C might be involved in mesendoderm specification (Madrigal et al. 2020). The identification of the DDIT3, IRF, JUN, and TFAP2 factors was consistent with the previous bulk RNA-seq results (**Fig. 1B**). However, the role of these markers in endoderm formation is not yet understood, although recent studies suggest that they might be involved in the process (Kojima et al. 2021; Li et al. 2019). For ME, we identified the following activated TFs: CDX1/2/4, which are involved in hematopoiesis (Paik et al. 2013), TCF4 (Kardon, Harfe, and Tabin 2003), LEF1 (Galceran et al. 2004), SP5 (Weidinger et al. 2005), HOXB (Iimura and Pourquié 2006), TBX6 (Sadahiro et al. 2018), and FOXH1 (David and Massagué 2018), which is also involved in heart development (von Both et al. 2004). The role of other top ranked TFs in ME development, such as MAX, ZNF90, DBP, E2F1, and TFDP2, remains unknown (**Fig. 2E**). SCENIC also assigned high activities to the pluripotency markers POU5F1 (OCT4), NANOG, and MYC in stem cells. SMADs were not among the top SCENIC hits, since their expression levels remained constant during the differentiation course, while SCENIC activities are based on the correlation of expression values of TFs and their target genes. Noteworthy, the output of SCENIC was not limited to TFs, for example, assigning a high activity to HDAC2 in stem cells compared to other stages.

Finally, we performed gene set enrichment analysis (GSEA) at the single-cell level, identifying high TGFβ and low WNT pathway activities in DE cells (**Fig. 2F**). Noteworthy, the activity of the mTORC1 Hallmark pathway was significantly reduced in DE cells compared to hESCs and PS. We also observed downregulation of the MYC pathway and activation of the hypoxia pathway in DE and ME compared to hESCs. GSEA of SMAD targets revealed a strong enrichment of SMAD2, 3 and 4 in PS and DE stages, corroborating their important role in DE differentiation. These results were consistent with those of the bulk RNA-seq data (**Fig. 1B**).

Taken together, scRNA-seq provided additional evidence for the importance of TGFβ/SMADs activation and mTOR/WNT inhibition in DE differentiation. It also uncovered new TFs that could be important for lineage specification at the PS branch point.

### Drug repurposing identifies candidate definitive endoderm inducers

We performed a CMap analysis using ssCMap (S.-D. Zhang and Gant 2009) on 300 common up- and down-regulated genes from the bulk RNA-seq analysis (150 for each cohort; **Methods**; **Fig. 3A**, right panel). The ssCMap tool relies on the initial release of CMap (with 6,100 conditions). Similarity of ssCMap profiles to the DE differentiation profile was assessed by GSEA (Subramanian et al. 2005). The analysis led to the identification of the PI3K inhibitors LY-294002 (henceforth LY29) and wortmannin, as well as the mTOR inhibitor sirolimus, whose expression profiles significantly correlated with the gene expression signature of DE (PC3 cell line) (**Fig. 3B**; see **Table S5** for the complete list). The utility of this approach is highlighted by the previous demonstrations that LY29 and wortmannin have been used along with AA to induce DE (Kuo, Liu, and Rajesh 2017; Pan and Liu 2019), and mTOR inhibition has been shown to improve mesendoderm induction (J. Zhou et al. 2009). Using the signatures of the same 300 common up-/down-regulated genes, we performed GSEA on the updated CMap catalog, LINCS. We set a threshold on the enrichment score to 0.35, resulting in the selection of 313 out of ~200,000 conditions corresponding to 298 unique compounds (**Fig. 3C**; **Table S5**). Interestingly, some of the top identified hits were heat shock protein 90 (HSP90) and histone deacetylase (HDAC) inhibitors. HSP90 has been reported to inhibit mesendodermal fate (Bradley et al. 2012), while HDAC inhibitors have been linked to endoderm lineage formation (Q.-J. Zhou et al. 2007; Rambhatla et al. 2003)(Goicoa et al. 2006). Together with ssCMAP, this analysis resulted in 299 (298 that include both sirolimus and wortmannin plus LY29) unique compounds. Later we refer to this approach as “ssCMAP/LINCS” (**Figs. 3B** and **C**).

**Fig. 3.**
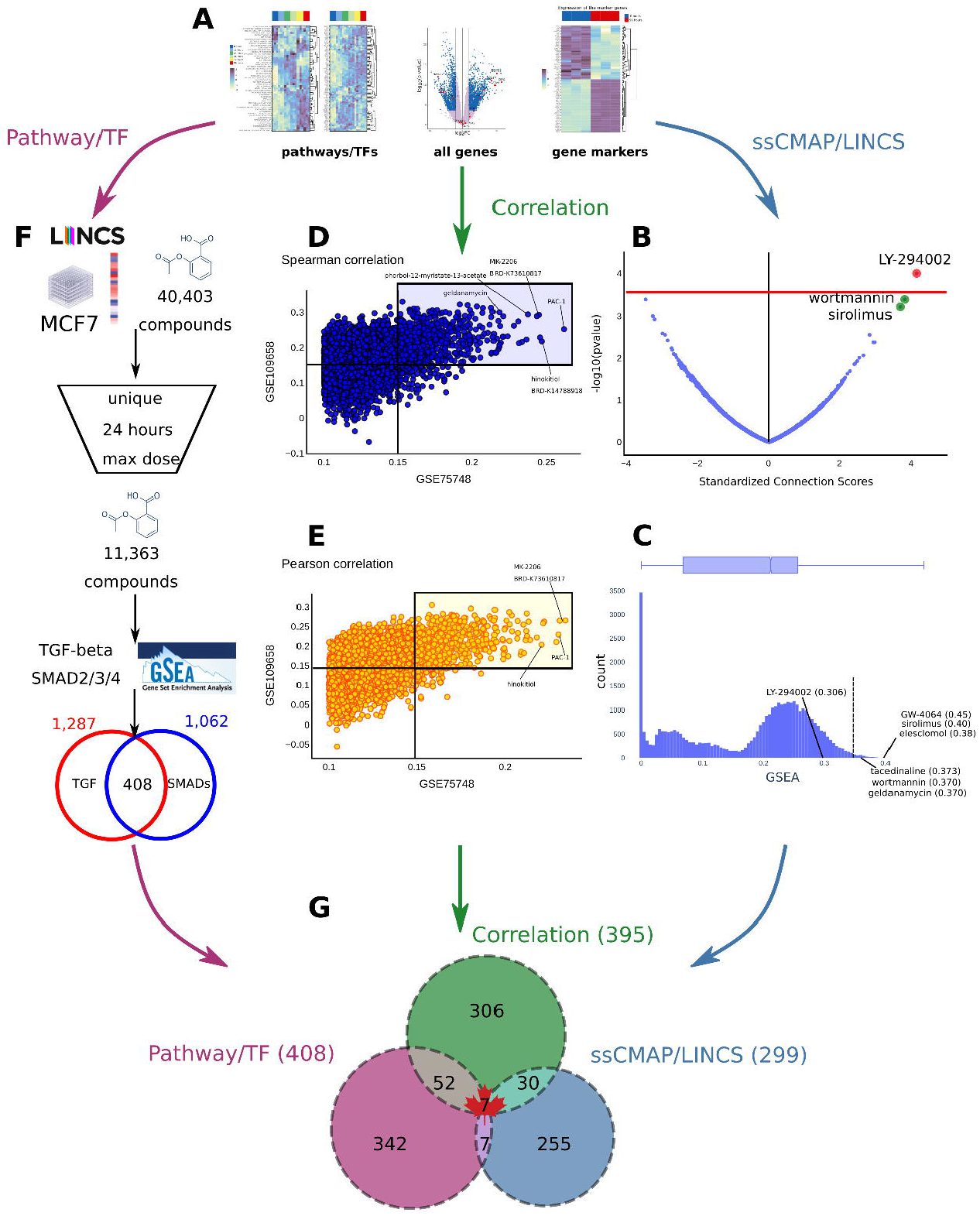
Drug repurposing pipeline. **A** Different sources of input for drug repurposing; all are based on the RNA-seq results of the public datasets; **B** - ssCMAP result for endoderm transcriptomic profile (intersection of GSE109658 and GSE75748); **C** - Distribution of top 40,000 compound signatures ranked by GSEA enrichment score. The threshold of 0.35 is highlighted with a dashed line; **D** and **E** - Spearman (D) and Pearson (E) correlation of LINCS signatures with transcriptomic profiles from the public datasets (GSE75748 - x axis; GSE109658 - y axis); correlation threshold of 0.15 for both datasets is highlighted with transparent rectangles; **F** - Pipeline for the identification of potential TGFβ/SMAD inducers. MCF-7 drug profiles are filtered based on dose and duration of treatment. The filtering is followed by GSEA with Hallmark and KEGG pathways and TF targets from RegNetwork; **G** - overlap between three different approaches;

As an alternative to using a subset of genes for GSEA in drug reference profiles, we compared the differential expression metric of all genes (*i.e*. the T-statistic) with the LINCS perturbation Z-scores using Pearson and Spearman correlations (**Fig. 3A**, middle panel). Concretely, we looked at how compound profiles from LINCS are correlated with the full gene expression profiles from both public RNA-seq datasets (GSE75748, GSE109658). This approach (henceforth “Correlation” approach (**Figs. 3D** and **E**)) has the advantage that it avoids thresholding (no fixed size gene sets). Both correlation metrics agreed well within and between datasets (**Figs. 3D** and **3E**; **Table S5**), and results were consistent with the “ssCMAP/LINCS” approach: we observed the presence of PI3K and mTOR inhibitors, including LY29, wortmannin, and sirolimus, as well as HDAC and HSP90 inhibitors, such as geldanamycin and tacedinaline. Setting a threshold on the Pearson and Spearman correlation of 0.15, we identified 395 unique compounds. The overlap with the 299 “ssCMAP/LINCS” compounds is 37 unique molecules.

During the bulk and single-cell RNA-seq data analyses, we observed TFs/pathways that could potentially induce DE differentiation (**Figs. 1B** and **3A**, left panel), including SMAD2, 3 and 4 known to be essential for the activation of the TGFβ signalling pathway (Nakao et al. 1997). Instead of looking at individual genes, alternatively, we searched for compounds that could activate the pathways and respective TFs. Using GSEA, we identified a list of molecules from LINCS whose profiles were enriched for TGFβ pathways, as well as for genes regulated by SMADs (**Fig. 3F**). This approach is based on comparing pathway/TFs enrichment profiles instead of individual genes. Therefore, it is “orthogonal” to the two previous pipelines. We refer to it as the “Pathway/TF approach”. Briefly, we focused on the cell line with the most abundant data in LINCS, MCF-7, which epigenetically resembles hESCs in terms of bivalency of its promoters (Messier et al. 2016). Prior to the analysis, all LINCS profiles were processed similarly to classical differential expression analysis (**Methods**). Overall, 1,287 compounds were enriched for the TGFβ signalling pathway according to definitions from either Hallmark (Liberzon et al. 2015) or KEGG pathways (Ogata et al. 1999). Similarly, 1,062 compounds were enriched for SMADs targets according to the RegNetwork database (Z.-P. Liu et al. 2015). As readers may be interested in other terms/pathways, the entire set of results is provided (**Table S6**). A total of 408 unique molecules overlap between both datasets (**Table S5**).

Taken together, we applied three drug repurposing strategies (**Fig. 3**): “ssCMAP/LINCS” approach with pre-defined gene markers (299 compounds); “Correlation” approach comparing entire DE differentiation profile to expression signatures of all compounds from LINCS (395 molecules); and “Pathway/TF” approach with LINCS compounds profiled only on MCF-7 cell line (408 molecules). The overall number of unique molecules was 999. Out of the 408 compounds from the last strategy, for practical reasons (*i.e*. cost and availability), we purchased 10 for testing. The reasons for purchasing compounds only from the third approach are detailed in the Discussion. The full set of results is provided in **Table S5.** The three strategies identified seven common compounds (**Fig. 3G**, **Table S5**), which were not experimentally validated but will be a priority in future research.

### Experimental validation of the compounds reveals new definitive endoderm inducers

In order to test candidate compounds for their ability to induce DE formation, we designed an endoderm reporter using CRISPR, as previously described (Krentz, Nian, and Lynn 2014): a SOX17-mNeonGreen (SOX17-mNG) knock-in H1 hESC line that preserved the endogenous *SOX17* mRNA sequence (**Fig. 4A**). Wild-type H1 hESCs and SOX17-mNG cells were differentiated into DE cells for three days (**Figs. 4B** to **4D**) using the standard protocol: 100ng/ml of AA + 3μM of CHIR for one day, followed by 100ng/ml of AA for two days (*i.e*. AC-A-A; **Fig. 4E**). No changes in the number of CXCR4+ cells were observed by FACS (**Fig. 5A**), suggesting that SOX17-mNG targeting did not impact endoderm differentiation.

**Fig. 4.**
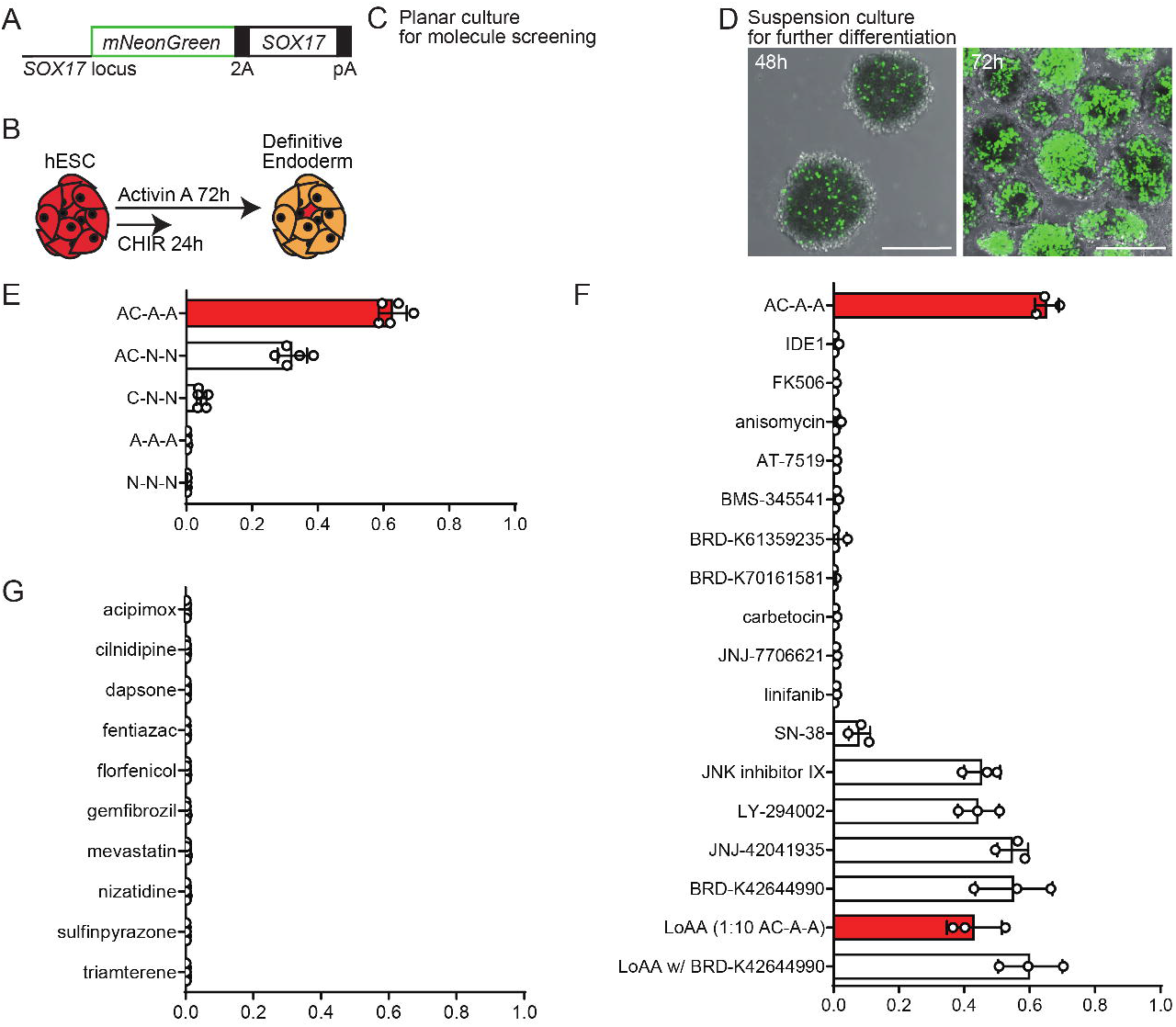
SOX17-NG system for molecule screening. **A** - Schematic representation of mNeonGreen cassette used in this study; **B** - Schematic representation of hESC differentiation toward DE with AA protocol; **C** - mNeonGreen-2A-SOX17 knock-in H1 (WA01) cells (homozygous mutant) differentiated by planar culture for 3 days; **D** - Suspension culture with standard protocol. Days 2 and 3; **E** - DE-induction efficiency oriented by AA. AC-A-A: Activin A and CHIR99021 on day 1, AA on day 2 and 3. AC-N-N: AA and CHIR on day 1, and no morphogens on day 2 and 3. etc; **F** - DE-induction efficiency by small molecules. Every treatment includes CHIR on day 1 and AA is replaced by each molecule described on Y-axis. LoAA: Treatment with CHIR on day 1 and AA by 1:10 concentration for 3 days. LoAA w/ BRD-K42644990: Treatment with CHIR on day 1, BRD and AA simultaneously by 1:10 concentration for 3 days; **G** - DE-induction efficiency by small molecules, MCF7 LINCS profile of which is not enriched for TGF-beta or SMADs targets. Every treatment includes CHIR on day 1 and AA is replaced by each molecule described on Y-axis.

**Fig. 5.**
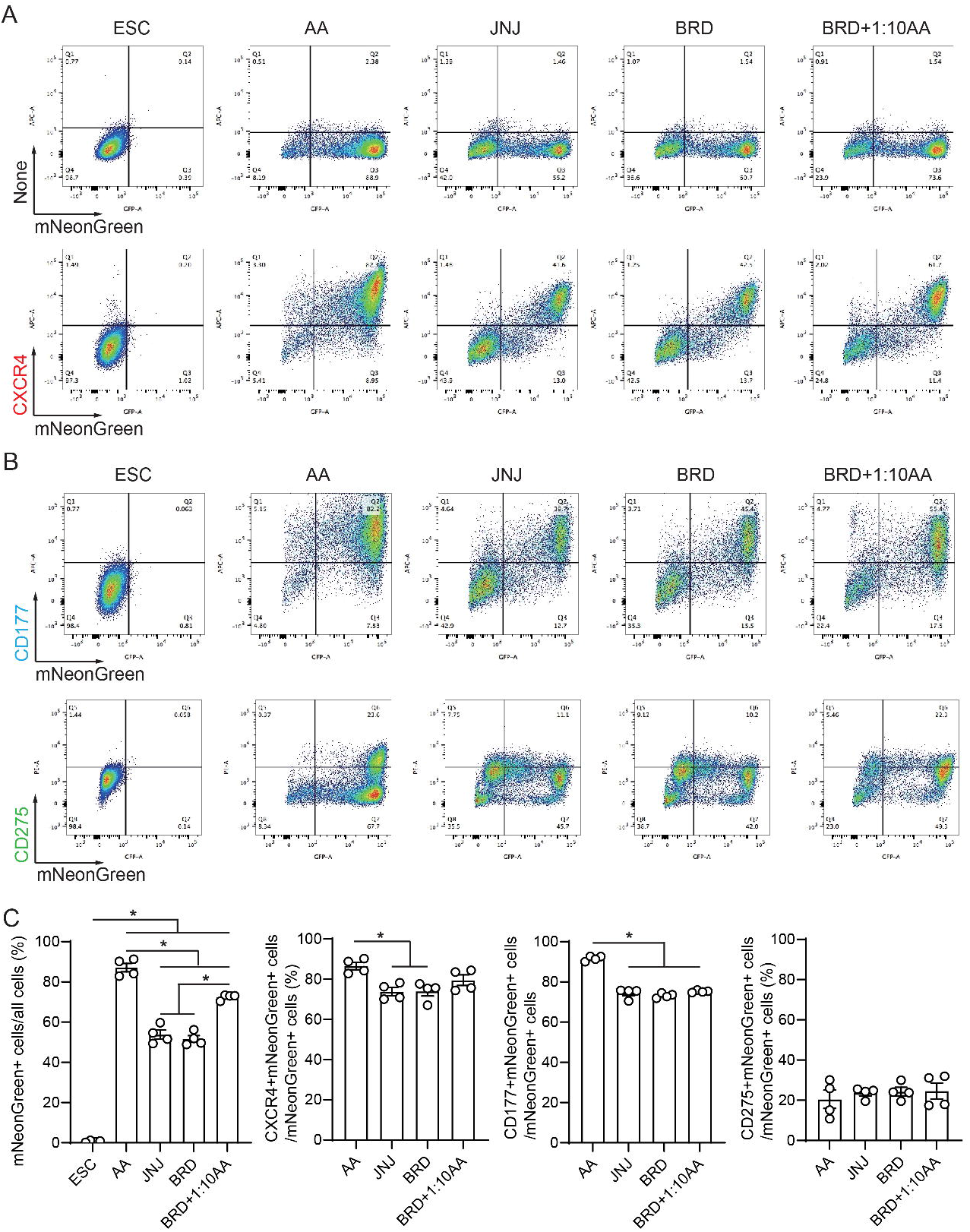
Characterization of DE cells induced by small molecules. **A** FACS data of DE cells differentiated from ESCs for 72 hours by each treatment. DE markers (CXCR4 and mNeonGreen reporter for SOX17) are analyzed. **B** FACS data of induced DE cells for surface markers CD177/CD275 indicating DE heterogeneity. **C** Quantification analysis of induced DE cells for mNeonGreen, CXCR4, CD177 and CD275. ESC: Non-treated ES cells as negative control. AA: Activin A and CHIR99021 on day 1, AA on day 2 and 3 as standard treatment. JNJ: JNJ and CHIR on day 1, and JNJ only treatment on day 2 and 3. BRD: BRD and CHIR on day 1, and BRD only treatment on day 2 and 3. BRD+1/10AA: Treatment with CHIR on day 1, BRD and AA simultaneously by 1:10 concentration for 3 days. \**P* < 0.05; n = 4.

High content screening (HCS) was conducted using a confocal high-content imaging system to quantify SOX17-mNG-expressing cells at various differentiation stages. On differentiation day 3, SOX17-mNG+ cells comprised 59.3 ± 2.1% of Hoescht 33342 staining cells (**Fig. 4E**), compared with 87.1 ± 2.1% using FACS (**Fig. 5A**). The lower endoderm induction efficiency observed with the HCS system was likely the result of being unable to exclude dead cells that would be Hoechst 33342+ and mNeonGreen- (**Figs. 4E**, **4F**, and **5A** to **C**). We next tested the importance of AA for DE induction. A single-day of AA+CHIR treatment induced 32.2 ± 2.0% cells to form DE (*i.e*. AC-N-N; **Fig. 4E**). Alone, neither CHIR (*i.e*. C-N-N) nor AA treatments (*i.e*. A-A-A) could induce DE formation (4.7 ± 0.7% and 0.2 ± 0.0%, respectively; **Fig. 4E**). These results confirm that the combination of CHIR and AA is required for inducing the formation of DE.

Next, we tested the 10 compounds selected based on the drug repurposing analysis for their capacity to induce DE in the absence of, or with a reduced concentration of AA (**Fig. 3F**; **Table S5**). Taking into account recent studies (Li et al. 2019; Madrigal et al. 2020), we were interested in testing a small molecule JNK inhibitor. As the 408 compounds from the pathway/TFs enrichment approach included several JNK inhibitors that were not accessible, we included an alternative JNK inhibitor that was available. LY29 was added to the compound list due to its strong drug repurposing support (““ssCMAP/LINCS” and “Correlation” approaches, see above) together with JNJ42041935 (henceforth JNJ), a HIF1 stabilizer that mimics a hypoxic condition. Hypoxic treatment of hESCs has been shown to enhance DE formation (Chu et al. 2016), suggesting that a physiological hypoxic condition during early development contributes to formation of DE cells. However, HIF1 stabilizers have not been used for *in vitro* DE formation. First, we determined whether the candidate compounds could improve DE differentiation after an initial one-day treatment with AA; eight out of twelve drug repurposing compounds (ten plus JNKi and LY29, ~70%) were more efficient than the baseline treatment (*i.e*. AC-N-N). Then, we sought to determine whether they could induce DE in the absence of AA; JNKi, LY29, BRD-K42644990 (henceforth BRD), and JNJ could (**Fig. 4F**). DE induction by BRD and JNJ was further confirmed by FACS (BRD, 51.7 ± 1.6%; JNJ, 53.8 ± 2.2%) (**Fig. 5B**). As negative controls, we screened 10 additional compounds that were not expected to activate TGFβ nor SMAD signaling pathways according to our third drug repurposing approach; none induced DE formation (**Fig. 4G**), confirming the robustness of the drug repurposing strategy used.

BRD is a novel compound with an unknown mechanism of action and, therefore, we sought to uncover its function in the context of DE induction from hESCs. To test whether BRD could replace AA, at least partly, regarding its cost effectiveness, we investigated its efficacy for DE induction by combining BRD with a reduced amount of AA. Treatment with 3 μM of CHIR for the first day only and BRD + 10 ng/mL of AA (1/10 of the concentration used in the standard protocol) for all three days induced 59.8 ± 4.6% of SOX17-mNG+ cells as detected by the HCS system, and 72.5 ± 0.7% as detected using FACS, representing a DE efficiency similar to that of the standard protocol but at a 90% cost reduction (**Fig. 4F**). Therefore, among the four compounds identified that were able to induce DE formation in the absence of AA, the subsequent analyses focused on JNJ and BRD (10 ng/mL AA).

### Population analysis of BRD-/JNJ-induced DE cells and subsequent differentiation into pancreatic endoderm

Similar to DE *in vivo, in vitro*-differentiated DE is heterogeneous: it contains CD177+ and CD275+ cell subpopulations with a preference for differentiating into either pancreatic or liver lineages, respectively. A recent study found that ~60% of DE-differentiated H1 hESCs were CD177+ (*i.e*. pancreatic lineage) and ~40% CD275+ (*i.e*. liver lineage) after a three-day treatment with AA (Mahaddalkar et al. 2020). To assess whether BRD/JNJ-induced DE cells achieved similar levels of heterogeneity, we performed FACS on differentiation day 3 (**Fig. 5B**). We found that mNeonGreen+ endodermal cells differentiated with BRD, JNJ and/or AA were predominantly CD177+, suggesting a preferential competence for pancreatic progenitor cells (**Fig. 5C**).

Next, we tested whether BRD and JNJ-induced DE cells could further differentiate into organ-specific cells such as pancreatic progenitors. Gene expression levels of pancreatic progenitor markers *NKX6-1* and *PDX1*, and the endocrine progenitor marker *NEUROD1*, were comparable between BRD-, JNJ-, and AA-induced DE cells (**Fig. 6A**). Moreover, for NKX6-1 and PDX1, protein expression levels were also comparable by flow cytometry (**Fig. 6B**). Finally, immunostaining revealed that NKX6-1 and PDX1 proteins were expressed in differentiated spheroids on day 15 (**Fig. 6C**). Taken together, these results suggest that BRD- and JNJ-based protocols can form endoderm that is largely similar to that generated with AA, and that BRD- and JNJ-induced DE can differentiate towards the pancreatic progenitor and endocrine stage with similar efficiency as with DE generated with AA.

**Fig. 6.**
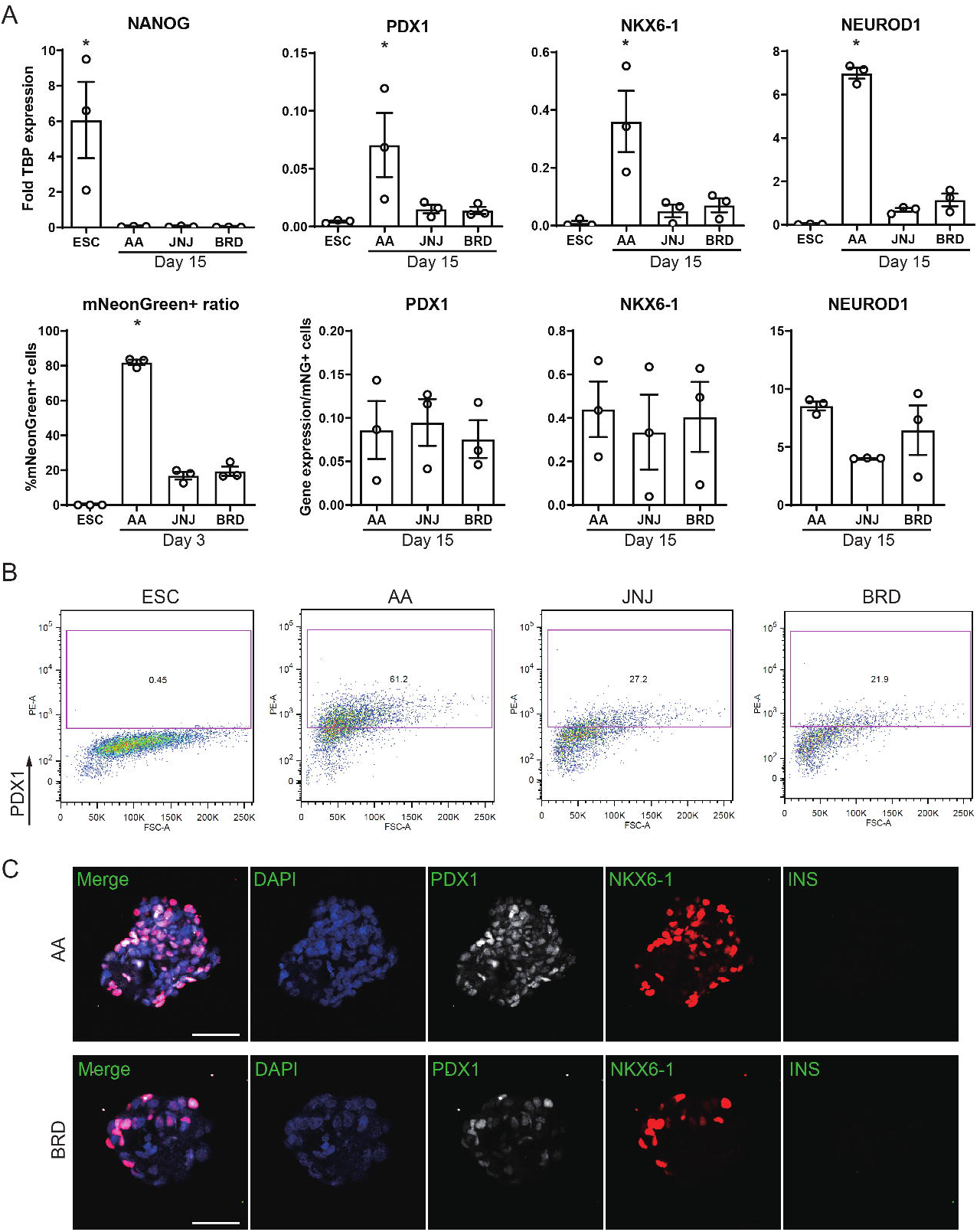
Characterization of pancreatic progenitor cells induced by small molecules. **A** mRNA expression levels of a stem cell marker (Nanog) and pancreatic progenitor markers (PDX1, NKX6-1 and NEUROD1) for pancreatic progenitor cells differentiated by small molecules for 15 days are quantified by nCounter assay (Nanostring) (upper panels). Each mRNA expression level is adjusted by the ratio of mNeonGreen cells in all cells on differentiation day 3 (lower panels). \**P* < 0.05 compared to the other three groups; n = 3. **B** PDX1 protein-positive cells are quantified for Day 15 PP cells by FACS. **C** Immunostaining for PDX1 and NKX6-1 proteins on AA- and BRD-induced PP cells. Scale bar, 50 μm.

### RNA-seq analysis of BRD-induced DE differentiation reveals downregulation of the MYC pathway

Both the targets and mechanism of action of BRD are unknown. Using DeepCodex (Donner, Kazmierczak, and Fortney 2018) and SwissTargetPrediction (Daina, Michielin, and Zoete 2019), which can predict a compound’s targets based on its similarity to known molecules, we did not obtain any significant hits. To explore its mechanism of action, we designed a bulk RNA-seq experiment wherein we tested the effect of a BRD treatment on gene expression alone and in combination with CHIR (BRD+CHIR), and compared it against a control pluripotent state and the standard AA protocol. To ensure that we would identify BRD-specific effects, we compared the BRD+CHIR and BRD-only treatments against a CHIR-only protocol (**Fig. 7**). Furthermore, we included a JNJ with CHIR (JNJ+CHIR) treatment as a positive control—another small molecule that can induce DE and with a known mechanism of action. Finally, since low doses of AA could increase the efficiency of the BRD+CHIR treatment, we also tested the sample from this modified treatment at 72h.

**Fig. 7.**
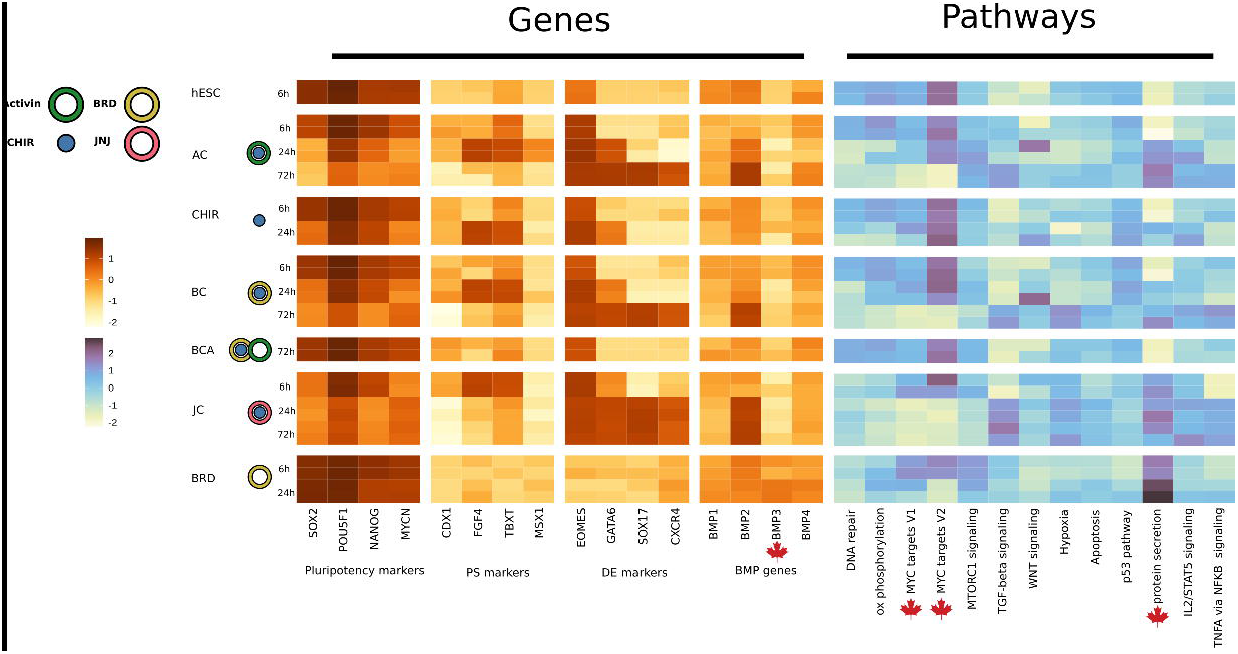
Effect of different compounds on hESC differentiation. Heat map of marker gene expression and pathway activities (Left) Scaled expression of the marker genes for different differentiation protocols; (Right) - single sample gene-set enrichment scores of selected Hallmark pathways.

An exploratory analysis of the bulk RNA-seq data revealed a strong confounding effect by CHIR during the initial differentiation stages; almost no genes showed significant differences in expression levels between the BRD+CHIR and CHIR-only treatments (**Table S7**). After 6h of BRD+CHIR treatment, we observed a significant increase in the expression levels of *EOMES* (vs. control pluripotent state), which is one of the key TFs guiding the endoderm lineage (**Table S7**) (Teo et al. 2011). However, AA induced additional regulators, including *MIXL1*, *TBXT*, *WNT3*, *CER1*, *FGF8*, and induced higher *EOMES* expression levels than BRD+CHIR (**Table S7**). Notably, AA strongly inhibited the pluripotency factor *SOX2*; in contrast, the expression levels of *SOX2* with both the BRD+CHIR and JNJ+CHIR treatments remained relatively high (**Table S7**). The expression profiles of key genes, such as *EOMES, SOX17*, and *GATA6*, for the BRD+CHIR and JNJ+CHIR treatments were almost identical, indicating either the existence of some commonalities or, most likely, a strong confounding effect by CHIR (**Fig. 7**). Interestingly, the only significantly expressed gene in the JNJ+CHIR vs. the BRD+CHIR treatment at 6h was *DDIT4*, a HIF1 responsive protein involved in mTORC1 inhibition (Brugarolas et al. 2004; Corradetti, Inoki, and Guan 2005) (**Table S7**). Finally, regardless of the treatment, we observed clear endoderm signals at 72h (**Fig. 7**; **Table S7**), however, although trends in expression levels were comparable, cells derived from the AA treatment displayed higher expression levels of endoderm-specific markers (**Fig. 7**).

Next, to characterize BRD-specific effects, we treated H1 hESCs with only BRD for 6 and 24h. The induced expression by BRD was clearly distinct from treatments leading to the endoderm lineage trajectory (**Fig. 7**). For instance, BRD induction of *EOMES* and other mesendoderm important genes were strongly downregulated, however, the expression levels of pluripotency markers (*SOX2*, *NANOG*, and *POU5F1*) were stable (**Fig. 7**). Noteworthy, the expression of *BMP3*, an inhibitor of both AA and BMP signaling (Gamer et al. 2005) that can activate SMAD2-dependent signaling by binding to ActRIIB (activin receptor type 2B) (Jialing Wen et al. 2019), was highly upregulated at 24h (**Fig. 7**, highlighted).

Shifting the focus of the analysis on the enrichment of gene terms, such as pathways and TF targets, we found that BRD-specific responses were mostly associated with down-regulated pathways, among which MYC was the most strongly inhibited both at 6 and 24h (**Fig. 7**, highlighted). Previous studies have found that a strong downregulation of MYC is correlated with high expression of meso- and endoderm markers (Cliff et al. 2017). The analysis also revealed a high activation of the protein secretion pathway, whose role in the differentiation process was unclear (**Fig. 7**, highlighted). Taken together, these results suggest a potential key role of MYC inhibition, and possibly BMP3 upregulation, in the mechanistic effect of BRD on pluripotent state exit and promotion of endoderm differentiation under WNT signaling via CHIR.

## Discussion

Using three orthogonal drug repurposing approaches, we identified a total of 999 unique candidates inducers for DE differentiation (**Table S5**). Exploration of the top hits indicates that these compounds are enriched for potential inducers: we observed the presence of PI3K, HDAC and mTOR inhibitors, some of which are used in existing protocols (Thakur et al. 2020), as well as HSP90 inhibitors, which are not currently used.

Exploring the three strategies, we prioritized the molecules found using our “Pathway/TF” approach. The reasoning was that while two other approaches were based on single gene expression levels, this one was focused on the activation of particular gene sets, which in theory is less noisy. Building a library and testing the other 989 compounds (if available) from across the approaches could be a focus of future research. Out of 408 unique “Pathway/TF” findings, we could purchase and process only 10 at the time. Notably, the 408 molecules included JNK inhibitors (such as JNK Inhibitor IX); JNK-JUN signaling has been recently shown to be a barrier towards endoderm differentiation (Li et al. 2019). Moreover, in a recent preprint the authors improved endoderm differentiation by using a JNK inhibitor (Madrigal et al. 2020). We wondered if it was possible to induce DE with a JNK inhibitor in the absence of AA. Therefore, we added one available JNK inhibitor to the purchase list, resulting in 11 compounds overall.

Overall, all three drug repurposing strategies did not agree broadly in terms of top compounds: “ssCMAP/LINCS” (299) and “Correlation” (395) overlapped with only 37 unique molecules, while each of them had in common with “Pathway/TF” strategy 14 and 59 compounds respectively. Even though these approaches are based on different methods (GSEA of the selected genes, correlation, and GSEA of specific gene sets) and data (in “Pathway/TF” approach we used only compounds that were profiled on MCF7 cell line), we still would expect a bigger overlap. These results may reflect limitations with both CMap- and LINCS-based drug repurposing approaches including the limited cellular context presented, approximation of gene expression values, inconsistency(Musa et al. 2017), and reproducibility (Lim and Pavlidis 2019). Thus, intersecting multiple methods and focusing on the limited subset of consensus compounds could be beneficial.

We tested the selected 11 candidate compounds for their ability to induce DE, as well as four previously described or novel endoderm inducers, including IDE1, LY29 (one of the CMap/LINCS findings and the supplement in some of AA-based DE differentiation protocols (McLean et al. 2007)), hypoxia inducer JNJ, and FK506BP inhibitor FK506i, making it 15 compounds total. When adding AA only for the first day, DE was induced by 11 out of 15 compounds. Among these, BRD, JNJ, JNKi, LY29, and FK506i were also able to induce DE formation in the absence of AA. Noteworthy, alone, IDE1 could not induce hESCs differentiation into endoderm, which is consistent with previous studies (Tahamtani et al. 2014). None of 10 randomly selected, negative control compounds tested could induce expression of endodermal markers. Previous blind screening approaches can also be considered as controls. For example, Korostylev *et al*. screened 23,406 compounds of which only 67 induced FOXA2 expression (endoderm marker) (Korostylev et al. 2017), and Borowiak *et al*. screened 4,000 compounds, identifying only two molecules, IDE1 and IDE2, that fit their criteria for DE induction (Borowiak et al. 2009).These results demonstrate that our *in silico* approach of candidate selection improves the outcome of screening studies. Moreover, we provide a valuable list of potential endoderm inducers that might be of interest to researchers planning a new drug screening procedure.

Among the identified DE inducers, BRD lacks any annotation, therefore, its targets are unknown. According to the LINCS pathway enrichment analysis, BRD is expected to be a TGFβ and SMAD inducer in MCF7 cells. However, our RNA-seq experiments revealed that this is not the case in hESCs. This highlights the limitations of drug repurposing approaches, as different cells often have distinct characteristics (Lim and Pavlidis 2019). In hESCs, BRD strongly inhibits the MYC pathway, while inducing the BMP3 (weakly) and protein secretion pathways. Both mechanisms seem relevant for DE differentiation: stauprimide, a molecule that primes hESCs toward DE is a known MYC inhibitor (Tahamtani et al. 2014); and BMP3 activates SMAD2 signaling (Jialing Wen et al. 2019). The relevance of protein secretion is unclear, however, the classical protocol with AA seems to induce the same pathway but to a lower extent (**Fig. 7**). Recently, replacement of AA by dorsomorphin, an inhibitor of the BMP ALK2 receptor, and titration of CHIR concentration have been shown to induce DE (Jiang et al. 2021). The authors also reported DE induction in the absence of AA, whereas another group observed no induction (Li et al. 2019), and our protocol only induced a small DE population (4.7%; **Fig. 4E**). These differences could be explained by different molecule compositions in each protocol, such as the existence of insulin and the absence of B27 in Li’s and our protocols, suggesting that the orchestration of multiple signaling pathways at various times is necessary for optimized DE induction. Of note, inhibition of BMP signaling was involved in DE induction both by dorsomorphin and BRD. Future experiments are needed for conclusive identification of BRD’s mechanism of action.

In addition to discovering AA replacements, we also generated a scRNA-seq dataset of the DE-ME branch. Our main goal was to discover key regulators of the branch point. Inhibiting the activity of lineage-specific TFs, especially in key developmental stages, might be a sound differentiation strategy (Graf and Enver 2009; Palii et al. 2019). For example, using a gradient of the TFs POU5F1 (OCT4) and SOX2 to control mesendoderm competence towards ectoderm fates (Valcourt et al. 2019). Therefore, it is tempting to speculate that a similar gradient of two or more TFs could be used to promote the transition of mesendoderm towards either DE or ME lineages. After performing clustering and differential expression analysis, we identified a set of TFs co-expressed at the PS branch point, but which then act as specific markers of either mesoderm or endoderm. The T-box TFs EOMES and Brachyury are among the top hits (see **Table S4** for the full list). Recent studies have demonstrated a potential cross-antagonizing effect between these TFs at a mesoderm-endoderm precursor stage (Tosic et al. 2019b; Chu et al. 2016). Brachyury is important for ME development (Faial et al. 2015), while EOMES is crucial for DE (Teo et al. 2011). Thus, inactivating Brachyury at this branch point with small molecules, or other factors, might promote differentiation towards endoderm (Faial et al. 2015; Tosic et al. 2019a). Aside from these TFs, other potential regulators might be of interest. For instance, inhibition of FOXH1, TBX6, SP5, and MSX1 will be a potential direction of future research to explore their effect on endoderm differentiation.

In conclusion, we have developed an *in silico* approach for identifying small molecules intended to optimize differentiation of pluripotent stem cells towards a variety of endodermal tissues. Our approach will be of utility to discover new molecules that could guide cell differentiation and reduce the expense of differentiation protocols.

## Methods

### Bulk RNA-seq analysis

Bulk RNA-seq data of hESC differentiation into DE were downloaded from GEO: accessions GSE75748 and GSE109658. For the GSE75748 dataset, which also includes single-cell RNA-seq data, the 0h time point was built using a pseudobulk approach by averaging individual cell counts from the single-cell data. Genes with <3 samples and >1 counts per million were filtered out from both datasets. For the in-house data, transcripts were quantified using salmon (1.4.0 version) (Patro et al. 2017). and annotated using GENCODE (version 36) (protein-coding transcripts only). Differential expression analysis was performed using the limma (3.48.1 version) (Ritchie et al. 2015) and edgeR (3.34.0 version) (Robinson, McCarthy, and Smyth 2010) R packages. A gene was considered as differentially expressed if the absolute change in log2-fold expression was >2, and significant (the FDR was controlled at 5% using the BH method).

### GSEA and heatmaps

GSEA was performed using the fgsea R package (1.18.0 version) (Sergushichev 2016). In the case of LINCS we performed the GSEA with marker genes using cTRAP R package (version 1.10.0) (de Almeida, Saraiva-Agostinho, and Barbosa-Morais 2021). The msigdbr R package (7.4.1 version) (Liberzon et al. 2015) was used to download the Hallmark and C5 (ontology) gene sets. The RegNetwork database (Z.-P. Liu et al. 2015) was used to identify TF targets. TF-gene connections were filtered for strong evidence only, and for TFs with literature support of directly binding to DNA (Lovering et al. 2020). A gene set was considered to be significantly up/down regulated if the normalized enrichment score was positive/negative respectively and higher than 1. The FDR was controlled at 5% using the BH method. For the single sample gene-set enrichment scores in bulk datasets the GSVA R package was used (1.40.1 version) (Hänzelmann, Castelo, and Guinney 2013). Pathway/TF activity heatmaps were plotted using the heatmaply R package (1.2.1 version) (Galili et al. 2018), and the clustering of pathways and TFs was based on gene expression data from the GSE75748 dataset. GSEA on scRNA-seq data was performed using the Escape R package (1.0.1 version) (Borcherding et al. 2021).

### Single-cell RNA-seq data

H1 hESCs were differentiated into DE cells using AA and CHIR99021 in a planar culture. Differentiated cells were collected at 36 and 72h post-induction as representatives of the sample including hESC and PS (36h) and the DE-enriched sample (72h). The mesoderm-cell sample was harvested at 72h post-induction with STEMdiff Mesoderm Induction Medium (Stemcell Technologies). Cells were grown on a 6-well plate. Prior to collection, they were washed once with PBS before adding 500 μL of Accutase. This was followed by 5 min incubation at 37°C for dissociation, after which we added 500 μL of 2% BSA in PBS, centrifuged the cells for 5 min at 200 × *g*, washed them once with PBS, and resuspended them in 350 μL of ice-cold PBS. scRNA-seq libraries were generated using a 10x Genomics ChromiumTM pipeline (Pleasanton, CA, USA), as previously described (Krentz et al. 2018). Single Cell 3’ Reagent Kits v2 were used. Briefly, cells were counted and loaded for a targeted cell recovery of 1000-5000 cells per channel. The microfluidics platform was used to barcode single cells using Gel Bead-In-Emulsions (GEMs). Reverse transcription was performed within GEMs, resulting in barcoded cDNA from single cells. The full length, barcoded cDNA was PCR-amplified, followed by enzymatic fragmentation and SPRI double-sided size selection for optimal cDNA size. End repair, A-tailing, Adaptor Ligation, and PCR were performed to generate the final libraries that have P5 and P7 primers compatible with Illumina sequencing. The libraries were pooled and sequenced using an Illumina NextSeq500 platform with a 150 cycle High Output v2 kit in paired-end format with 26 bp Read 1, 8 bp I5 Index, and 85 bp Read 2. Following sequencing, FASTQ files were generated from the raw sequencing data using cellranger mkfastq (10x Genomics), after which they were mapped to the build 38 of the Genome Reference Consortium human genome (GRCh38) to generate single-cell gene counts using cellranger count (10x Genomics). Low quality score sequences were filtered out. Cellranger aggr (10x Genomics) was used to combine data from multiple samples and ensure that all libraries had the same sequencing depth.

### Seurat analysis

The Seurat R package (4.0.2 version), executed with default parameters unless otherwise specified, was used for quality control, filtering, data preprocessing, scaling and normalization of scRNA-seq raw data. High quality cells with mitochondrial gene content lower than 5%, feature total number higher than 200 and lower than 4000 and RNA count number higher than 2500 were extracted. The top 5000 most variable genes were selected based on their average expression and dispersion, and used to reduce the dimensionality of the data through principal component (PC) analysis. The top 50 PCs were selected for clustering. tSNE (REF) were applied to the selected PCs for visualization of the data. Cluster-specific markers were identified using the Seurat’s FindAllMarkers function. Differentially expressed genes between two clusters were identified using the Seurat’s FindMarkers function with both the parameters min.pct and logfc.threshold set to 0.2.

### SCENIC analysis

pySCENIC (0.11.2 version), executed with default parameters (*SCENICprotocol: A Scalable SCENIC Workflow for Single-Cell Gene Regulatory Network Analysis n.d*.), was applied on the pre-processed scRNA-seq data to identify potential important TFs and their targets. First, potential modules of TFs and targets were identified based on their co-expression using GRNBoost2 (Moerman et al. 2019). Then, each co-expression module underwent *cis*-regulatory motif analysis using RcisTarget to filter out indirect targets without motif support and retain modules with significant motif enrichment. These RcisTarget-processed modules are referred to as “regulons”. Next, the activity of each regulon in each cell was computed using AUCell (Aibar et al. 2017). The resulting AUC scores were further processed using SCopeLoomR (0.11.0 version) (*SCopeLoomR: R Package* (*compatible with SCope*) *to Create Generic .loom Files and Extend Them with Other Data E.g.: SCENIC Regulons*, *Seurat Clusters and Markers*, n.d.) and scFunctions (0.0.0.9 version) (Wuennemann n.d.) for downstream analysis and visualization.

### Pseudotime analysis

Single-cell trajectory analysis was performed using the Monocle R package (2.9.0 version) using the output from the Seurat analysis as input. Monocle’s DDRTree method was applied with default parameters for dimensionality reduction to two dimensions.

### LINCS analysis

The LINCS analysis was performed as in Lim and Pavlidis (Lim and Pavlidis 2019). Samples with the same dose, molecule, time point, RNA plate, and cell line properties were regarded as the same experiment. Accordingly, conditions were matched with the appropriate control (same cell line and time point) to calculate the log2-fold change in expression for every gene. Since GSEA takes as input a list of ranked genes, we used log2-fold expression changes as a measure of “gene significance”.

### Drug repurposing

#### ssCMAP and LINCS with marker genes (GSEA)

Out of 735 common significantly up-regulated genes in the GSE75748 and GSE109658 datasets, we selected the top 150 for CMAP and GSEA analysis. Similarly, out of 433 common significantly down-regulated genes, we selected the bottom 150, resulting in 300 genes overall. The number of 150 up- and down-regulated genes was chosen based on default values of cTRAP R package (version 1.10.0) (de Almeida, Saraiva-Agostinho, and Barbosa-Morais 2021) and suggestions from CMap web service as a number that gives optimal results. Gene names were converted to microchip probe identifiers using the hgu133a.db R package (version 3.13.0) (Carlson 2021), resulting in a list of 343 probes. The list was used as input to ssCMap (S.-D. Zhang and Gant 2009), which aims to find a significant connection between the provided gene signatures and 6100 core reference profiles summarizing properties and activities of ~1000 small molecules. The 300 gene names were further used to perform GSEA on the LINCS cellular perturbation data using the cTRAP R package. Potential DE inducers (*i.e*. small molecules output from cTRAP) were filtered based on their enrichment (GSEA) score (>= 0.35) and adjusted *p*-value (<0.05).

#### Pearson and Spearman correlation of the gene expression

The T-statistics (from the limma analysis) of all genes in the GSE75748 and GSE109658 datasets were used to build two profiles, one for each dataset. The cTRAP R package was used to find compounds from LINCS that were positively correlated with these profiles. For robustness of the results, we used both the Pearson and Spearman correlation metrics. The most promising compounds were those with a correlation >0.15 in both datasets. Combination of results from the Pearson (584) and Spearman (371) analyses resulted in 395 unique compounds.

#### Pathway/TF enrichment and intersection

Compound log-fold gene expression signatures were selected from the MCF7 cell line, which covered the largest number of conditions (40,403) encompassing a total of 11,418 unique molecules. 24h of treatment and highest dosage were used as filters, resulting in 11,363 unique compounds (and 14,260 conditions). Each of the resulting signatures was used as input for GSEA with the following gene sets: Hallmark and KEGG pathways, and RegNetwork TF targets sets. We selected those compounds that were enriched for the TGF-beta pathway according to at least one annotation (either KEGG, 654 compounds, or Hallmark, 767 compounds). This filtering step resulted in 1,287 unique compounds. After this, we selected compounds that were enriched for the targets of SMAD2 (38), SMAD3 (739), and/or SMAD4 (718), resulting in 1,062 unique compounds. Finally, we intersected the two compound sets (pathways and TFs), which resulted in 408 unique compounds.

### Generation of SOX17-mNeonGreen hESC line

The SOX17-mNeonGreen hESC line was generated as described in Krentz *et al*. (Krentz, Nian, and Lynn 2014). Briefly, the CRISPR/Cas vector was based on px458 (Addgene; plasmid 48138)), exchanging the Cbh promoter for a full-length CAGGS promoter in order to maximize expression in hESCs (*i.e*. pCCC). The gRNA (AGAGTGGTGACGGAGACAGG; score 0.6) was designed using the algorithm from Doench *et al*. (Doench et al. 2014), and was cloned into the BbsI sites of pCCC to generate pCCC-LL801/2, as described in Ran *et al*. (Ran et al. 2013). The targeting vector was obtained from Addgene (plasmid pJet 1.2). The mNeonGreen-2A-SOX17 donor plasmid and pCCC-LL801/2 encoding plasmid were transduced into H1 hESCs by electroporation using Bio-Rad Gene Pulser II system. Cells were selected by 0.25 mg/mL puromycin (Sigma). Colonies were manually picked and cultured. Genomic DNA was extracted using QuickExtract (Epicentre) and the following primers were used for genotyping: 5’F CAGGCAAGTTGAGTCCTGGG, 5’R GTAACCGTCGTTTGGGTTGC, 3’F TTTTGGTCTCCTGGAAGCGG, 3’R ACCAGCCGATGTACGTGTTT. A homozygous mutant was used for experiments.

### Maintenance and differentiation of hES cells

hESCs were maintained on diluted Geltrex-coated (ThermoFisher Scientific; 1:100 in DMEM/F12) plates in StemFlex (Invitrogen). Cells were split every three or four days and plated at a density of 1×10^6^ per 60 mm plate. 1 mM Y-27632 dihydrochloride (Tocris Bioscience) was added for the day of split to keep cell viability. hESCs were differentiated using the protocol published by Nair *et al*. (Nair et al. 2019). For planar culture (Protocol 1), hESCs were plated onto Geltrex-coated 12-well plates at a density of 5×10^5^ in StemFlex medium with 1 mM Y-27632. Differentiations began 24h post-seeding. Cells were treated with RPMI (Gibco) containing 0.2% FBS, 1:5,000 ITS (Gibco), 100 ng/ml Activin A and 3 μM CHIR99021 on day 1. On day 2-3, cells were treated with RPMI containing 0.2% FBS, 1:2,000 ITS and 100 ng/ml Activin A. For suspension culture (Protocol 2), dissociated H1 hESCs were plated on 6-well plates at a of density of 5.5 million cells in 5.5 mL media per well. The plates were incubated at 37°C and 5% CO_2_ on an orbital shaker at 100 rpm to induce spheroid formation. After 24h, a 6-step differentiation was induced. Planar culture enabled the detection of mNeonGreen+ cells after 3 day-culture for the HCS of candidate DE-inducing molecules. Suspension culture allowed more efficient differentiation toward pancreatic progenitors by maintaining 3D structure. Media compositions were as follows: Day 1: RPMI (Gibco) containing 0.2% FBS, 1:5,000 ITS (Gibco), 100 ng/ml Activin A and 3 μM CHIR99021. Day 2: RPMI containing 0.2% FBS, 1:2,000 ITS and 100 ng/ml Activin A. Day 3: RPMI containing 0.2% FBS, 1:1,000 ITS, 25 ng/ml KGF (R&D Systems), 2.5 μM TGFbi IV (CalBioChem). Day 4–5: RPMI containing 0.4% FBS, 1:1,000 ITS and 25 ng/ml KGF. Day 6–7: DMEM (Gibco) with 25 mM glucose containing 1:100 B27 (Gibco), 3 nM TTNPB (Sigma) and 0.5 mM ascorbic acid. Day 8: DMEM with 25 mM glucose containing 1:100 B27, 3 nM TTNPB, 0.5 mM ascorbic acid and 50 ng/ml EGF (R&D Systems). Day 9–10: DMEM with 25 mM glucose containing 1:100 B27, 50 ng/ml EGF, 50 ng/ml KGF and 0.5 mM ascorbic acid. Day 11–15: DMEM with 25 mM glucose containing 1% fatty acid-free BSA, 1:100 Glutamax (Gibco), 1:100 NEAA (Gibco), 1 mM Pyruvate (Gibco), 1:100 ITS, 10 μg/ml of heparin sulfate, 1 mM N-acetyl cysteine (Sigma-Aldrich), 10 μM zinc sulfate (Sigma-Aldrich), 1.75 μL 2-mercaptoethanol, 10 μM Repsox (ALK5i, Stem Cell Technologies), 2 μM T3 (Sigma-Aldrich), 500 nM LDN-193189 (Stemgent), 1 μm Xxi (Millipore). Differentiation is stopped at day 15 to assess pancreatic progenitor cell state. Morphogens were replaced for specific experiments as described in the manuscript. On differentiation day 3, there was a difference in DE induction efficiency between the standard AA protocol (~90%) and the BRD/JNJ protocol (~60%) assessed by FACS. As a result, we FACS sorted the mNeonGreen+ cells on day 3 and reaggregated the cells to reestablish the 3D culture prior to continuing the differentiation processes. Unfortunately, we were unsuccessful at maintaining a reaggregated endoderm past day 4. Therefore, in an effort to control for endoderm induction efficiency, we normalized gene expression and protein levels detected on day 15 to the starting endodermal cell input.

### High content imaging system

ImageXpress Micro Confocal System (Molecular Devices) was used as a high content imaging system. For the HCS, a Corning 96-well plate was coated with Geltrex and incubated overnight. hESCs were plated with a density of 5×10^4^ cells per well and cultured in StemFlex (Invitrogen) with 1 mM Y-27632. After 24h, 3-day differentiation was induced. For nuclear staining, cells were incubated with Hoechst33342 at 37 °C for 10 min.

### Flow cytometry

Spheroids were collected and incubated with Accumax (Stem Cell Technologies, BC, Canada) at 37°C for 10 minutes, and dissociated into single cells for FACS. CMRL (Corning, NY) with 1% BSA was added, followed by filtering through a 40-mm nylon filter. Cells were centrifuged for 5 min at 200 × *g*, washed with PBS, and resuspended in 500 μL of PBS. Cells were incubated with APC mouse anti-Human CD184 (CXCR4) (BD), APC anti-human CD177 (REAfinity, Miltenyi Biotec), PE anti-human CD275 (B7-H2) (REAfinity, Miltenyi Biotec), or PE mouse anti-PDX-1 Clone 658A5 (BD) antibodies at room temperature for 30 min, followed by washing with PBS and resuspension in 500 μL of PBS. LSRFortessa (BD Bioscience, CA) was used for FACS.

### Gene expression assay

Sample cells were collected and incubated in the RLT buffer with 1% 2-mercaptoethanol. nCounter gene expression assay (Nanostring, WA) was performed to assess NANOG, PDX-1, NKX6-1, and NEUROD1 gene expression levels in hESCs, differentiated DE, and pancreatic progenitor cells using 1.5 μL of the samples. The values were normalized by TBP.

### Immunohistochemistry

For immunostaining, spheroids were fixed in 4% formaldehyde, 2% agarose gel-embedded, paraffin-embedded, and sectioned at 5 μm. Primary antibodies were applied overnight at 4°C in PBS-0.3% Triton+ 5% horse serum. After washing with PBS, secondary antibodies and DAPI were applied for 1h at room temperature, before mounting on slides using SlowFade Diamond Antifade Mountant (Invitrogen). Primary antibodies were rabbit anti-PDX1 (D8O4R) 1:400 (Cell Signaling, #54551), goat anti-NKX6.1 1:400 (Abcam, #AF5857) and guinea pig anti-INS 1:1000 (DAKO, #A0564). Secondary antibodies were donkey anti-guinea pig FITC, donkey, donkey anti-goat Cy3 and donkey anti-rabbit APC 1:200 (Jackson Immuno Research Laboratories, #706-096-148, #705-166-147, #711-136-152). Spheroids and sections were imaged using a Leica SP8 confocal microscope with a 20x/0.75 IMM objective.

### Statistical analyses for stem cell experiments

Measurements were performed on discrete samples unless otherwise specified. Statistical analyses were performed using the GraphPad Prism 8.0. Multiple groups were analysed by one-way ANOVA with a multiple comparison test and the Tukey-Kramer’s post-hoc test was used to compare different groups. A *p*-value < 0.05 was considered to indicate a statistically significant difference between two groups. Data are presented as the mean ± SEM.

## Code for generating the results

All relevant scripts are available on GitHub: https://github.com/fransilvion/Drug-repurposing-for-stem-cells.

## Acknowledgments

G.N. was supported by an International Doctoral Fellowship from the University of British Columbia. O.F., and W.W.W. were supported by grants from the Canadian Institutes of Health Research (PJT-162120), Natural Sciences and Engineering Research Council of Canada (NSERC) Discovery Grant (RGPIN-2017-06824), and the BC Children’s Hospital Foundation and Research Institute. F.C.L. was supported by the Canadian Institutes of Health Research (PJT-156377) and the Stem Cell Network. Salary (F.C.L.) was supported by the Michael Smith Foundation for Health Research (#5238 BIOM) and the BC Children’s Hospital Research Institute.

N.L. and P.P. were supported by NIH grant MH111099. N.L. was supported by UBC Four-Year Doctoral Fellowship.

Fellowship support was provided by the Juvenile Diabetes Research Foundation (S.S.; 3-PDF-2018-587-A-N), the Michael Smith Foundation for Health Research (S.S.; 17045), the Manpei Suzuki Diabetes Foundation (S.S.), The University of British Columbia (G.N.; International Doctoral Fellowship), and the CIHR (H.H.; CGS-M).

The authors acknowledge that UBC and BC Children’s Hospital are situated on the traditional, ancestral and unceded territories of the Coast Salish peoples – the S<u>k</u>w<u>x</u>wú7mesh (Squamish), Sǝlílwǝta?/Selilwitulh (Tsleil-Waututh) and x^w^mǝθk^w^ǝýǝm (Musqueam) Nations.

## Abbreviations

AA: Activin A
BRD: BRD-K42644990
CHIR: CHIR99021
CMap: Connectivity Map
DE: definitive endoderm
EMT: epithelial-mesenchymal transition
GSEA: gene set enrichment analysis
HCS: high content screening
HDAC: histone deacetylase
hESCs: human embryonic stem cells
JNJ: JNJ42041935
LINCS: Library of Integrated Network-based Cellular Signatures
LY29: LY-294002
ME: mesoderm
mNG: mNeonGreen
MsigDB: Molecular Signatures Database
PS: primitive streak
ssCMap: statistically significant Cmap
TF: transcription factor

---

**Fig. S1. Overview of the object of the study. A** - schematic representation of human embryonic stem cell (hESC) differentiation process towards definitive endoderm through primitive streak stage (a common precursor of endoderm and mesoderm cells); **B** Mechanism of action of Activin A; (left) before the binding of Activin A to a type 2 receptor, the cell is in “silent” mode: Alk4 cannot activate the TGFβ pathway through SMAD proteins since it is blocked by FKBP12; (right) upon binding, Activin A induces the interaction of a type 2 receptor with Alk4, which induces the removal of FKBP12 and phosphorylation of SMAD2/3. Phosphorylated SMAD2/3 forms a complex with SMAD4, which translocates to the nucleus and induces the expression of target genes, thereby activating the TGFβ pathway. Created with BioRender (“BioRender” n.d.).

**Fig. S2. Differential expression analysis of the public datasets. A** - PCA plot of the definitive endoderm differentiation protocol (GSE75748 time-series data). Arrows show the progression along the differentiation time axis; **B** - transcriptomic changes between two data sets (GSE75748 and GSE109658) are significantly positively correlated (Pearson correlation 0.57); **C** - Volcano plot of the 4-day differentiation time point compared to a pluripotent state demonstrates strong transcriptomic effect (GSE75748 left, GSE109658 right); key marker genes are highlighted in red circles.

**Fig. S3. Pathway and TF activities of GSE109658 dataset.** Heatmaps with activation values of pathways (left) and transcription factors (right) along differentiation timeline.

**Table S1.** Gene markers of human embryonic stem cells, primitive streak, definitive endoderm and mesoderm.

**Table S2.** Selection of genes for CMAP analysis. Commonly expressed genes - list of common up-regulated (735) and down-regulated (433) genes in GSE75748 (left) and GSE109658 (right) datasets; DGE statistics is shown for each gene; Drug_repurposing_genes - 300 genes that were selected as ssCMAP input; ssCMAP_input - 300 genes converted to AffyProbeSetID (one gene can have more than one probe ID); regulation column shows if a gene is up- or down-regulated.

**Table S3.** GSVA enrichment scores for pathways and TFs. top_pathways_75748, top_pathways_109658 - enrichment scores of Hallmark pathways for GSE75748 and GSE109658 public datasets respectively with the statistic; top_TFs_75748, top_TFs_109658 - enrichment scores of TF targets (RegNetwork) for GSE75748 and GSE109658 public datasets respectively with the statistic; top_pathways_109658;

**Table S4.** Differentially_expressed_TFs - differential expression statistic for detectable TFs in different states vs hESCs; RSS_(SCENIC)_scores_of_each_cell_type - regulon specificity scores for different regulons for each of the cell type;

**Table S5.** top_GSEA_compounds - compounds that pass the enrichment score threshold of 0.35; best_Pearson_Spearman_compounds - compounds, gene expression profiles of which are correlated with a DE differentiation profile higher than 0.15 Pearson and Spearman correlation score (for both GSE75748 and GSE109658); MCF7_TGF_SMAD_inducers - list of compounds that either induce TGFβ pathway or SMAD2/3/4 TFs (according to the enrichment analysis). Some of the compounds are repeated, they correspond to different RNA plates; purchased compounds - list of purchased compounds that were enriched for TGF and SMADs TFs at the same time; common_compounds - 7 small molecules that were found by all three drug repurposing approaches

**Table S6.** The activity of different compounds in terms of Hallmark and KEGG pathways, and TF targets. 1 - positive enrichment; −1 - negative enrichment; 0 - not significant;

**Table S7.** The results of differential gene expression analysis and Hallmark pathway GSEA comparing BRD differentiation conditions to other protocols;

