## Supplementary figures and images for "*In silico* discovery of small molecules for efficient stem cell differentiation into definitive endoderm"

### Sfig1_corAA.png

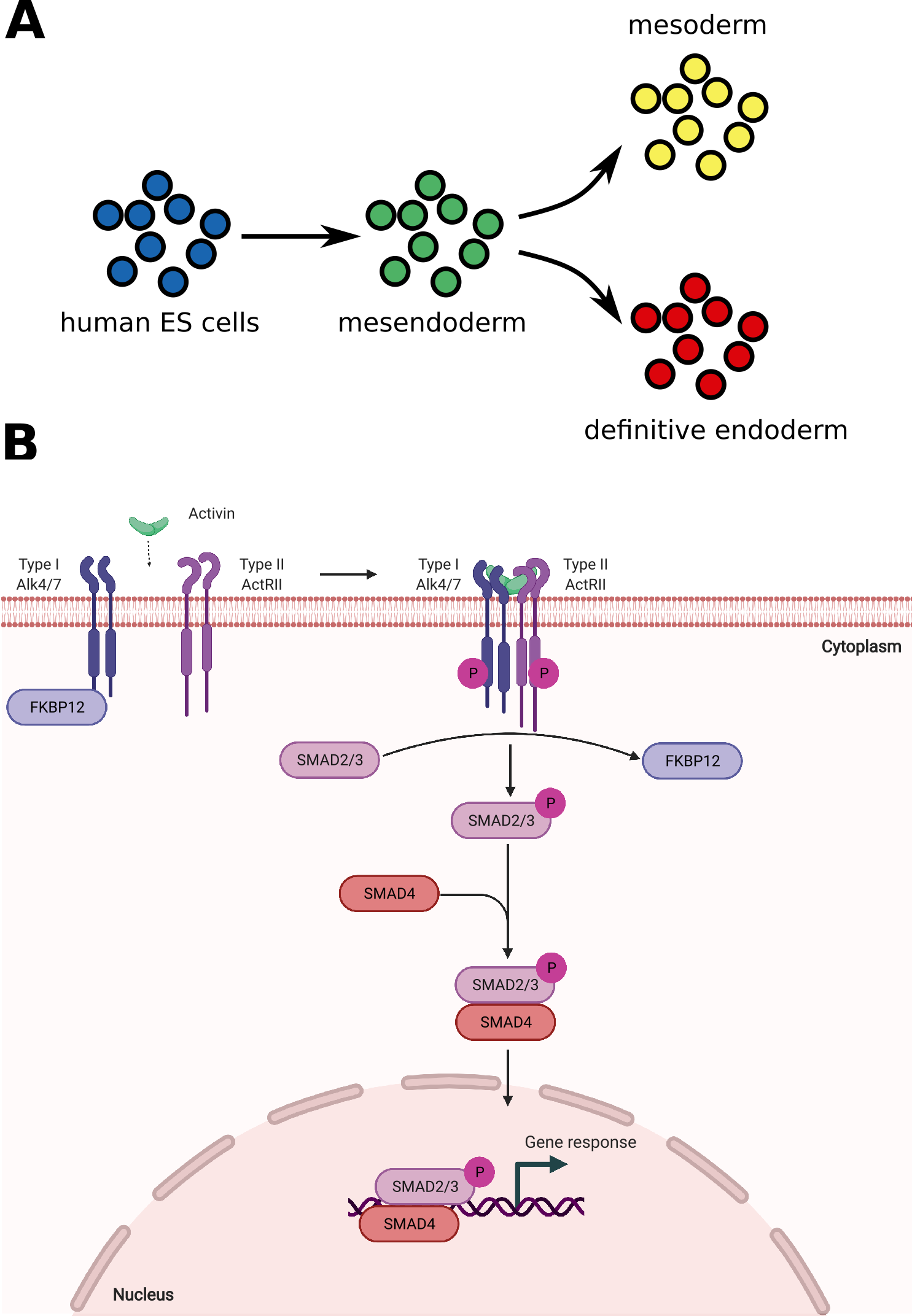

### Sfig2.png

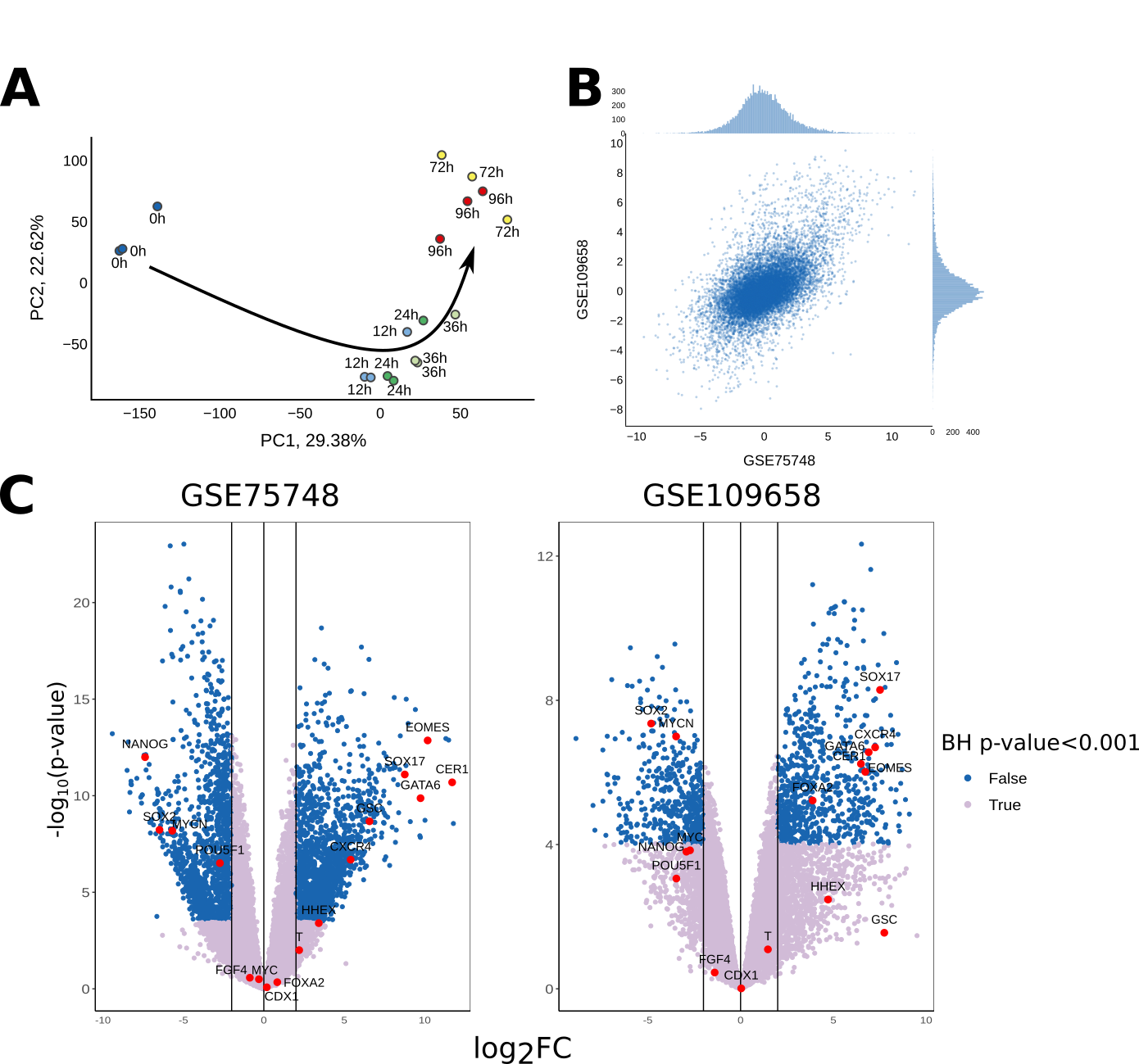

### Sfig3.png

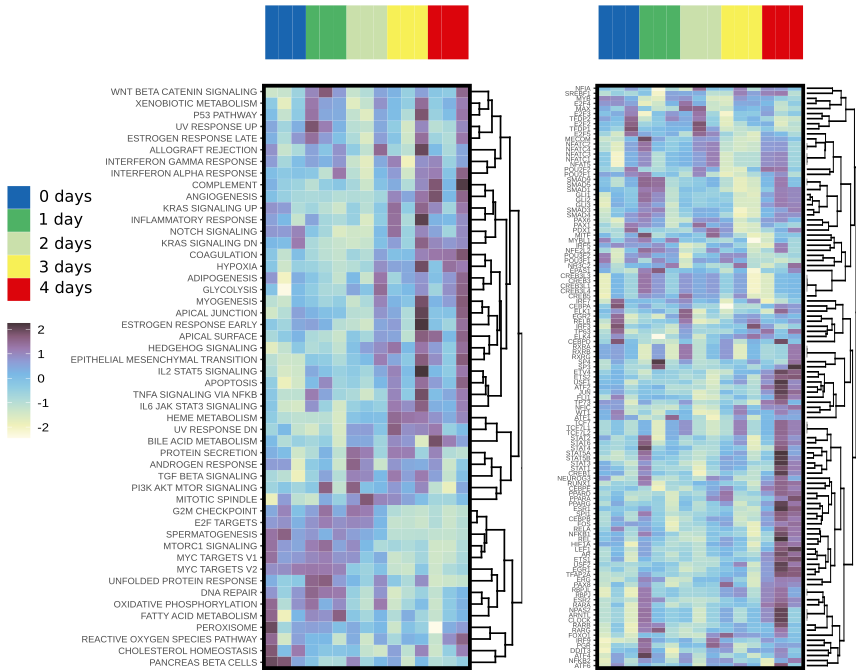
